# Design of intrinsically disordered region binding proteins

**DOI:** 10.1101/2024.07.15.603480

**Authors:** Kejia Wu, Hanlun Jiang, Derrick R. Hicks, Caixuan Liu, Edin Muratspahić, Theresa A. Ramelot, Yuexuan Liu, Kerrie McNally, Sebastian Kenny, Andrei Mihut, Amit Gaur, Brian Coventry, Wei Chen, Asim K. Bera, Alex Kang, Stacey Gerben, Mila Ya-Lan Lamb, Analisa Murray, Xinting Li, Madison A. Kennedy, Wei Yang, Zihao Song, Gudrun Schober, Stuart M. Brierley, John O’Neill, Michael H. Gelb, Gaetano T. Montelione, Emmanuel Derivery, David Baker

**Affiliations:** Department of Biochemistry, University of Washington, Seattle, WA, USA; Institute for Protein Design, University of Washington, Seattle, WA, USA; Biological Physics, Structure and Design Graduate Program, University of Washington, Seattle, WA, USA; Department of Electrical Engineering and Computer Science, University of California, Berkeley, CA, USA; Department of Chemistry and Chemical Biology, Center for Biotechnology and Interdisciplinary Studies, Rensselaer Polytechnic Institute, Troy, NY, USA; Department of Chemistry, University of Washington, Seattle, WA, USA; MRC Laboratory of Molecular Biology, Cambridge, CB2 0QH, UK; Visceral Pain Research Group, Hopwood Centre for Neurobiology, Lifelong Health Theme, South Australian Health and Medical Research Institute (SAHMRI), North Terrace, Adelaide, South Australia 5000, Australia; Faculty of Health and Medical Sciences, University of Adelaide, North Terrace, Adelaide, South Australia 5000, Australia; Howard Hughes Medical Institute, University of Washington, Seattle, WA, USA

## Abstract

Intrinsically disordered proteins and peptides play key roles in biology, but the lack of defined structures and the high variability in sequence and conformational preferences has made targeting such systems challenging. We describe a general approach for designing proteins that bind intrinsically disordered protein regions in diverse extended conformations with side chains fitting into complementary binding pockets. We used the approach to design binders for 39 highly diverse unstructured targets and obtain designs with pM to 100 nM affinities in 34 cases, testing ∼22 designs per target (including polar targets). The designs function in cells and as detection reagents, and are specific for their intended targets in all-by-all binding experiments. Our approach is a major step towards a general solution to the intrinsically disordered protein and peptide recognition problem.

## Main

Natural evolution has generated a variety of solutions to the challenge of binding unstructured regions of peptides and intrinsically disordered proteins (IDPs)(*1–6*), including natural antibodies(*1–3*), the major histocompatibility complexes(*4*), tetratricopeptide repeats(*5*), Armadillo repeat proteins(*6*), and lipocalins (Anticalins). Despite this diversity, engineering general peptide recognition remains challenging: peptide-specific antibodies have been obtained by immunization or by library selection, but this requires considerable effort, and disordered antigens are susceptible to degradation following injection. There has been progress in generalizing the binding modes of armadillo repeat and other natural peptide-binding proteins(*3–5*), but achieving completely new specificities has been challenging. De novo protein design methods have been used to design proteins that bind peptides in polyproline II, alpha-helical and beta-strand conformations(*7–9*), but more general recognition of disordered proteins and peptide regions requires the ability to bind more varied conformations as an arbitrary disordered sequence may not have propensity for the same secondary structure throughout, or present suitable interfaces for binding in regular secondary structures. For example, amphipathic helices or strands can be recognized using designs with grooves that bind primarily to the non-polar side of the helix or strand, but if charged residues are distributed around the helix or strand axis, this binding mode would require energetically unfavorable charge burial. A method for achieving specific recognition of any target intrinsically disordered region (IDR) sequence of interest would be broadly useful for applications in proteomics, targeting, sensing, and sequencing.

We reasoned that a general solution to the IDR binding problem might be achieved by combining the strengths of physical and deep learning design approaches. Rosetta design methods have been used to design binding proteins consisting of four to six tandemly repeated structural units that bind repeating proline-rich sequences in the polyproline II conformation, with each repeat unit in the designed binder interacting with a repeat unit in the peptide (Fig. 1a). Generative deep learning free diffusion methods(*8*, *10*) have been used to generate binding proteins with no such pre-imposed structural constraints on the binder structure (Fig. 1b), but because the model is trained on the PDB, this procedure generally folds the target sequence into the alpha-helical or beta-sheet conformations that dominate the PDB, which as noted above can be poorly compatible with binding. The repeat protein-repeat peptide approach can, in principle, be generalized beyond polyproline II conformations, but is limited by the requirement that each peptide unit have the same conformation, which again may be incompatible with heterogeneous target sequences. We reasoned that starting from different repeat protein architectures and recombining and specializing amino acid binding pockets in different repeat units for different amino acids and different conformations using diffusion could yield a family of templates enabling more general recognition of sequences with widely varying conformational preferences and sequences (Fig. 1c). In the following sections, we first describe the creation of such a scaffold library for general peptide recognition (Fig. 1d-f), and then the use of the library to design binders for a wide variety of non-repeating, both synthetic and native IDR targets (Fig. 1g-j).

**Fig. 1.**
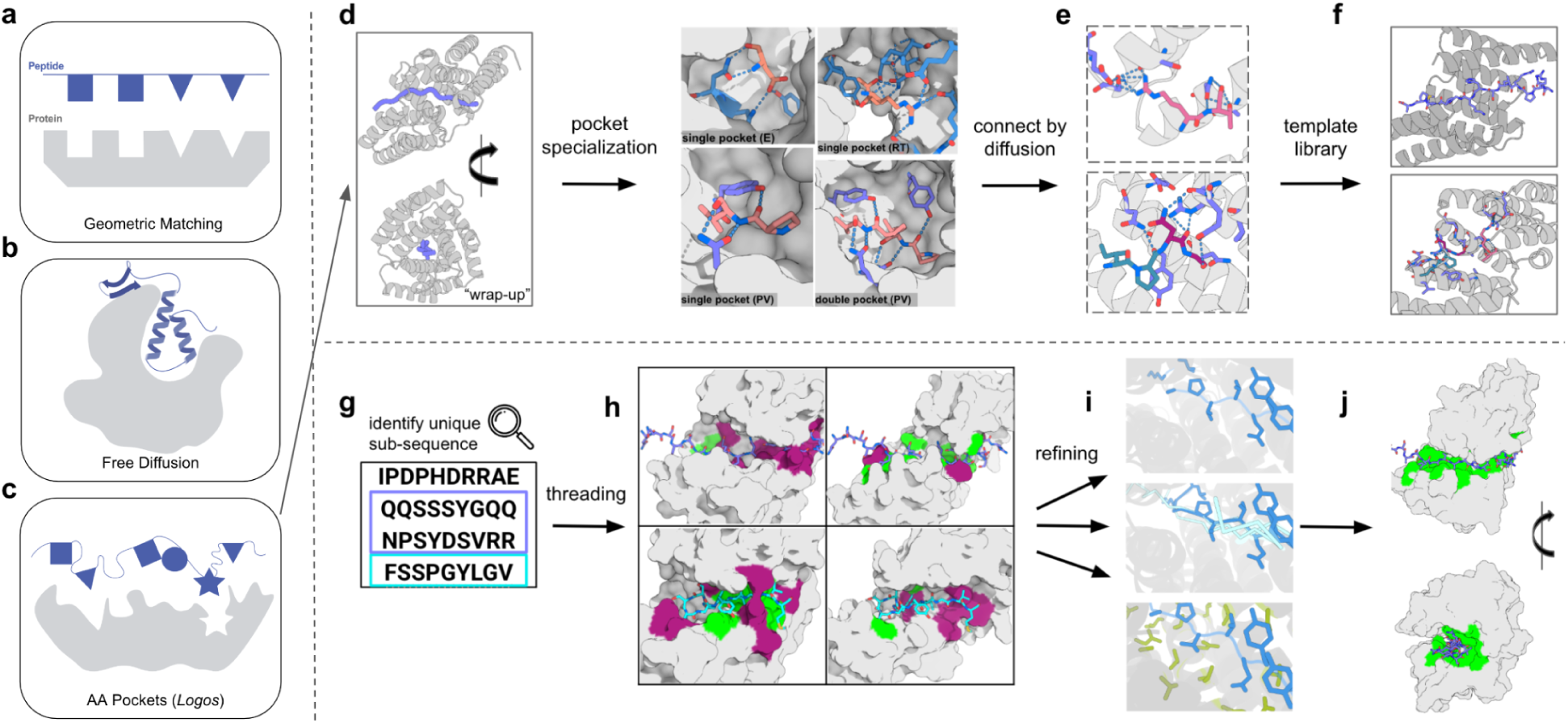
Overview of IDR binder design protocol. **a-c, Design methods. a,** Repeat protein based geometric matching approach requires one-to-one matching between identically spaced repeat units on designed binder and target peptide . **b,** Unconstrainted free diffusion approach folds targets into structures frequently observed in PDB training set, primarily helices and strands. **c,** The AA pockets approach explored here combines the designed pockets and extended scaffolds of the geometric matching approach with the ability of diffusion to recombine and diversify the pockets to achieve more general recognition of non-repeating sequences. **d-f, Template library construction. d,** Left, designed binder scaffolds wrap around extended peptide backbones, enabling contacts with each target amino acid. Right, example binding pockets. **e,** Binding pockets are connected using RFdiffusion (each peptide window colored differently) into templates for general sequence recognition. **f,** Examples of two of the 1000 generated templates. **g-j, IDR binding pipeline. g,** Unique sub-sequences (purple and cyan) are identified through a protein sequence database search, and **h,** threaded through the template library to identify optimal matches between amino acid segments and binder pockets. Pocket matches are green and mismatches are dark red on the protein surface. **i,** Matches are refined using “one-sided partial diffusion” (top), where only the binder is changed; “two-sided partial diffusion” (middle), where the target and the binder can be changed; “motif diffusion” (bottom), where key interacting motifs (target, blue; binder, green) are unchanged while the rest are noised, reconnected, diversified, and optimized. **j,** Examples of resulting designs.

### Generation of Template Library

We reasoned that the template library should have two properties. First, each template structure should “wrap” around extended peptide conformations with numerous opportunities for the hydrogen bonding and packing interactions with the target required for high specificity (Fig. 1d). Second, the structural variation in the template family should be sufficiently broad that for any target sequence at least one of the templates is able to induce it into a defined binding-competent conformation.

We developed a three step approach to generating such a library of template structures suitable for general recognition. In the first “scaffold generation” step (Fig. 1d; Methods I), we design repeat proteins that wrap around peptides in different repeating conformations, such that each repeat unit on the protein forms a binding pocket that interacts with a corresponding repeat unit on the peptide. We require that these pockets have side chains that not only interact with the target side chains but also make hydrogen bonds with the target backbone to provide structural specificity and compensate for the cost of desolvation. In the second “pocket specialization” step (Fig. 1e; Methods II), we fine-tune these pockets using diffusion to achieve more precise matching to specific target peptide sequences. We keep the four to nine amino acids surrounding each sidechain bidentate hydrogen bond from the repeat protein to the peptide backbone fixed (Fig. S1) while diversifying the hydrophobic interactions between designed binder; this is advantageous because hydrogen bonding interactions have more stringent geometric requirements than non-polar packing interactions and hence are more efficiently templated than repeatedly sampled from scratch (see Methods II-4). In the third “pocket assembly” step, we go beyond the limitations of repeating structures, which are optimal for repeating sequences but not more general sequence targets, by recombining pockets from different designs, using RFdiffusion(*11*) to generate interfaces between them where necessary to yield overall rigid structures (Fig. 1f). This generates a set of templates with diverse pockets arranged in different orders and geometries (Fig. 1f; Methods III).

For the first “scaffold generation” step, we chose to target peptides in a broad range of extended conformations rather than solely the polyproline II conformation as in our earlier study, as this is populated primarily by proline-rich peptides. In extended conformations, alternating side chains face in opposite directions, consistent with a two-residue sequence repeat. We used Rosetta design methods as described previously to generate designs targeting the dipeptide repeats LK, RT, YD, PV, and GA (single-letter amino acid codes) in a variety of extended conformations that wrap around these peptides such that each repeat unit interacts with one dipeptide unit (Fig. 2a; Methods I). Experimental characterization by fluorescence polarization of four-repeat versions of the designed binders revealed nanomolar binding for the LK and PV repeat peptides, but little binding for the more polar RT and highly flexible GA (Fig. S2; to avoid potentially unfavorable interactions with peptide termini, for experimental testing here and below, we pad all repeat target peptides with two additional repeats).

**Fig. 2.**
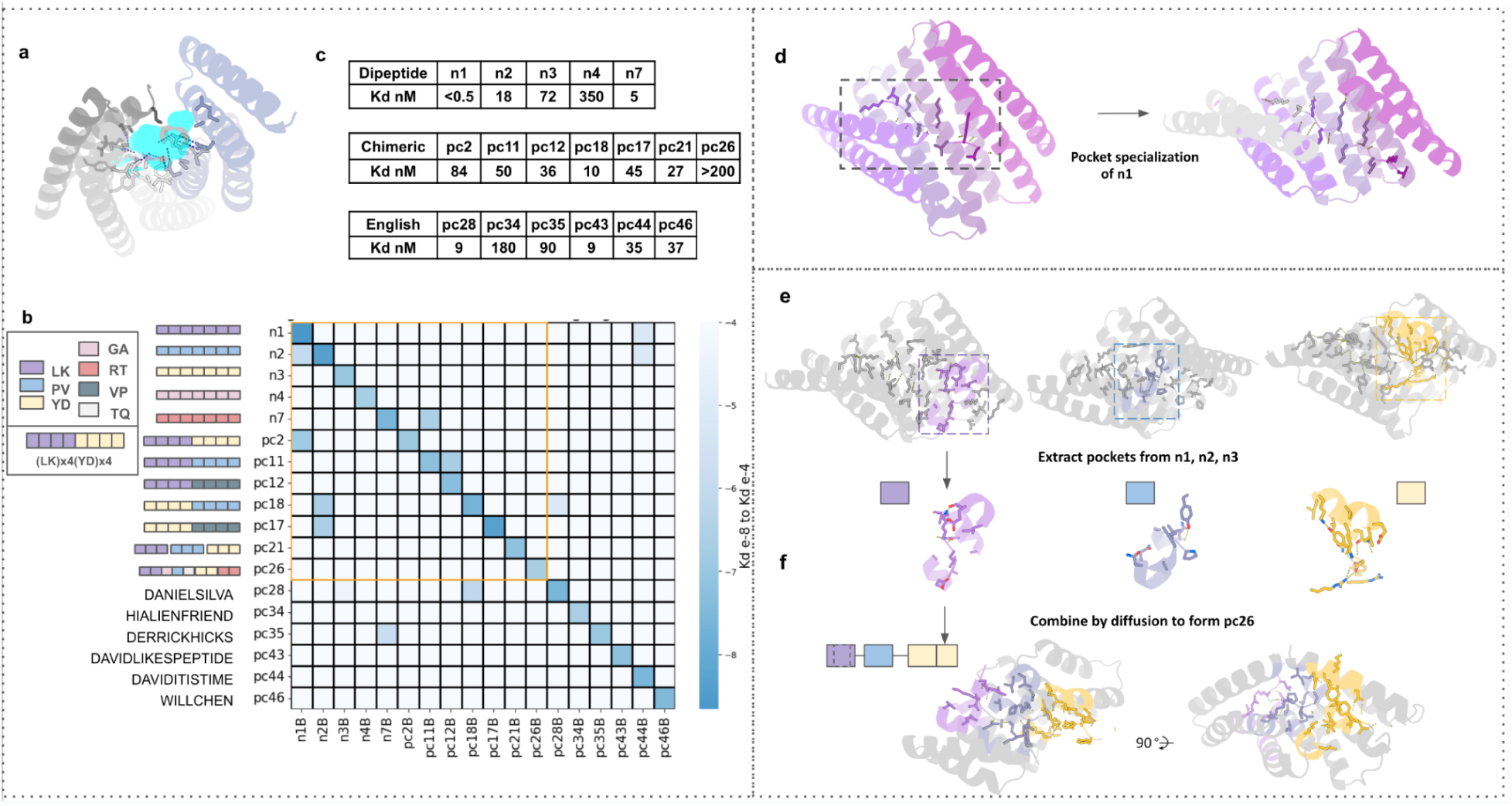
Design of binding to 18 synthetic peptides. **a,** Representative binder design (cartoon) wrapping around peptide (cyan) in an extended conformation (hydrogen bonds shown as dotted lines). **b,** All-by-all binding Kd’s obtained from nanoBiT binding titrations for 18 designed binder-synthetic peptide pairs. Target sequences are on the y-axis with each square representing one dipeptide motif using the color scheme in the legend (left). Heatmap intensities indicate Kd averages from N=2 titration experiments. Pairs within the orange square are composed of similar dipeptide repeats and hence have more cross-talk. **c,** Cognate designed binder-target Kd’s measured by biolayer interferometry. Peptide label-identity pairings are on the y axis in b. Two to 35 binder designs were experimentally tested per target. **d-f,** Library construction example using pc26. The three recombined pockets are shown using color codes from panel **b**. **d,** “Pocket specialization,” the repetitive pockets are optimized and specialized using motif diffusion from the old four-repeat scaffold (left) to the new five-repeat scaffold (right). The extended fifth repeat is shown in light grey. **e,** Extracting pockets containing two-sided interacting motifs from the base dipeptide repeats. **f,** Diffusion flexibly connects multi-motifs into coherent binding proteins with varied spacers and angles. An example of an assembled design is shown at the bottom.

For the second “pocket specialization” step, we refined the designed binding pockets to improve the fit to the target sequences and extended the number of interacting repeat units from four to five to further increase affinity. This yielded designs with picomolar affinities for LK repeats, and low nanomolar affinities for RT and GA repeats (Fig. 2b-c; Fig. S3). The binding pockets in these designs have distinct geometries that are customized to the target being recognized (see example in Fig. S4).

For the third “pocket assembly” step, we enable more general recognition of non-repeating sequences by assembling the binding pockets into new backbones, keeping them positioned to interact with peptide targets in continuous extended conformations (Fig. 2d-e). We assembled combinations of two to six binding pockets in silico (see below and Methods III), yielding models of chimeric proteins interacting with chimeric peptide targets. To do this, we positioned pockets parametrically (see Methods III-1), and connected them with RFdiffusion. We used this approach to generate 70 designs against seven chimeric targets. We refer to each binding unit (comprising a single amino acid or dipeptide and corresponding designed protein pocket) with a letter; thus, AAABBB is a chimera of two designs from the previous section, while ABCDEF combines six different pockets. Experimental characterization using nanoBiT split luciferase reconstitution(*12*) and biolayer interferometry (BLI) showed double-digit nanomolar binding for six out of seven of the targets, out of only ten designs tested per target on average (Fig. 2b; Fig. S4).

To increase the size of the template library to cover a broader range of sequences, we used pocket assembly to build 36 chimeric backbones containing pockets recognizing polar residues and further diversified both binder and peptide target by two-sided sequence design in silico (see Methods III-3; in the designs described above, the peptide sequence was always held constant). Together, this yielded a library of 1,000 templates, each consisting of a designed binding protein and a corresponding peptide backbone positioned such that the amino acids in the peptide fit into designed pockets in the binding protein (representative examples shown in Fig. S5).

### Threading Intrinsically Disordered Regions onto Template Library

We developed a two step approach for using the template library to generate binders for non-repeating synthetic sequences and arbitrary native unstructured targets. In the first “threading” step (Fig. 1g), the target sequence is threaded through the backbone of each template to identify the most compatible sequence segment-template pairs. In the second “refinement” step, the best matches are refined to increase the fit between the designed binder and target peptide (Fig. 1h-i).

For an IDP or IDR, there are, in general, a large number of possible peptide subsequences that can be targeted. To identify the most targetable peptide subsequences within an IDR, we first discard segments with low sequence complexity and/or with multiple close matches in the proteome (Fig. 1g, S6, and S7), as binders to such targets would likely have some cross-reactivity. We map each of the remaining unique sequence segments of eight to 40 amino acids onto each of the target backbones in the library, carry out local backbone resampling, optimize the sequence of the binder using ProteinMPNN, and evaluate the designs based on the fit between the designed binder and the target sequence and the agreement between the AF2 prediction and the design model (see Methods IV). This approach maps target segments with multiple polar residues into templates compatible with extended hydrogen bonding networks, which is likely important for achieving general recognition. In cases where AF2 metrics were suboptimal, we used RFdiffusion (see Methods V) to customize the backbone for the specific target.

We first tested this approach on synthetic sequences corresponding to six arbitrarily selected English words and names. We tested 45 designs against six targets, eight designs per target on average; the best binders for two out of the six targets had single-digit nanomolar affinities (K_d_s of 9 nM), three had double-digit affinities (K_d_ = 35 nM, 37 nM, 90 nM), and one a K_d_ = 180 nM (Fig. 2b; Fig. S8). We investigated the selectivity of the designs for their peptide targets by carrying out all-by-all (18 by 18) nanoBiT interaction measurements, including the repeat sequence binders of the previous section. Although there was some cross-talk between designs and targets with related sequences (for example, designs targeting four PV repeats also bound peptides with eight PV repeats), for the more diverse targets, the designs were quite orthogonal (Fig. 2c; Table S1).

We next used the threading approach to generate binders for 21 diverse therapeutically relevant IDPs, IDRs, and segments of IDPs ranging from eight to 40 amino acids. These include eight GPCR ligands, two insulin-related ligands, four disease detection-related disordered regions, four IDRs from cancer-related receptors, and three human scaffolding complexes for which there are no good monoclonal antibodies (target names and targeted sequences in Fig. 3a). For each target, three to 48 designs (on average 28) for which the AF2 predicted structures of the complexes were close to the computational design models (Fig. S8) were selected for experimental characterization. The designs have overall helical architectures reflecting their helical repeat protein origins (Fig. S9), with considerable structural variation in some regions introduced by the diffusion assembly and refinement steps, while the target sequences adopt a wide range of random coil and partially helical and strand conformations (Fig. 3b-c; Fig. S10).

**Fig. 3.**
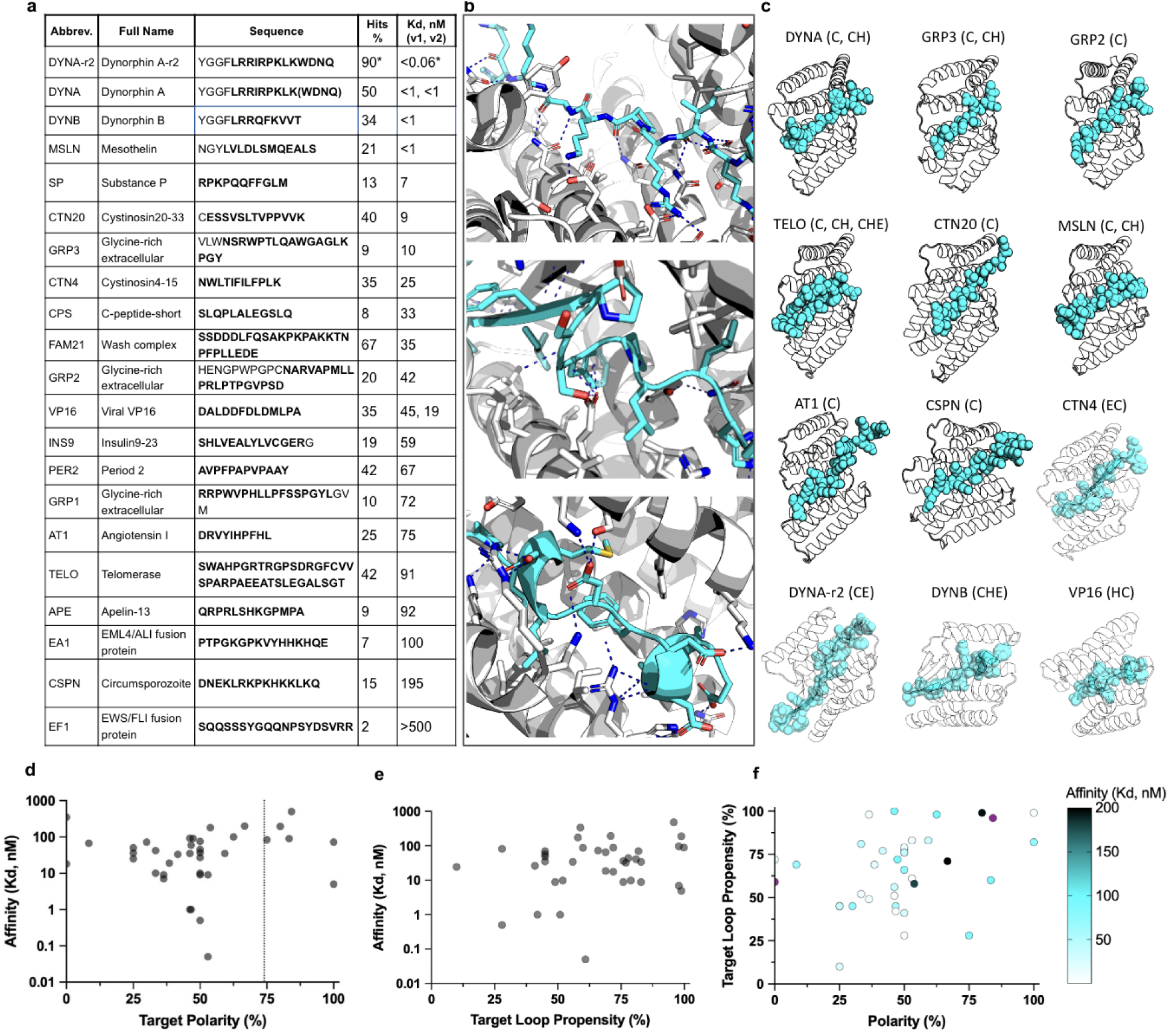
Design of binding to 21 native protein disordered regions and peptides. **a,** Targeted native bioactive peptides and IDRs; bold indicate the interacting sequences. Best obtained Kds are ranked from low to high; for the same target, the best two Kds from binders targeting two distinct target conformations are separated by commas. Hit rate was calculated based on the percentage of tested designs showing binding signals (> 0.1 AU) on BLI at 500nM. Asterisks indicate statistics calculated with the optimization campaign, instead of a one-shot campaign. Three to 48 designs were experimentally tested per target. **b,** Interactions of representative IDR binder conformations: random coil conformation (top); strand-containing conformation (middle); helical-containing conformation (bottom). **c,** Models of representative designed complexes. Abbreviations of the targets are above each model, with a bracket indicating the designed targeting conformations of the target. C = random coils, H = (partial) helix, E = (partial) strand. Commas separate multiple targeting conformations for the same target. **d,** Target polarity vs. highest achieved affinities. Polarity calculated based on the percentage of polar and charged amino acids among the targeted window. **e,** Target loop propensity calculated with AIUPred3 algorithms vs. highest achieved affinities. **f,** Target polarity vs. target loop propensity with a color gradient showing the affinities (black, outliers; dark blue, 200 nM; white, low pM).

The designed binders were expressed and purified, and binding to the targeted IDRs was measured by BLI with the peptide attached to the sensor chip and the binder in solution. The fraction of binders that showed a binding signal at 500 nM ranged from 2-67% (Fig. 3a). Together with the 18 synthetic targets, we obtained binders for 39 of 43 targets, testing on average 22 designs for each target (Table S2).

Polar targets have been long considered challenging in protein design. Twenty of the targets had more than 50% polar residues, and six had more than 75%; binders were obtained for the fusion fragment EF1 of EWS/FLI onco-fusion protein for Ewing sarcoma(*13–15*) (84% polar residues), and the N-terminal fusion fragment CSP-N of Circumsporozoite protein (CSP) for malaria(*16*) (80% polar residues) (see Fig. 3d for target polarity distribution). The number of hydrogen bonds to the side chains of the target (per 10-amino-acid segment) was three-fold higher for the highly polar targets than the other targets (4.3 for target polarity <75%; 12.3 for target polarity >=75%). 77% of the targets had little predicted secondary structural propensity in isolation; 87% adopted predicted bound conformations lacking extensive secondary structure, and all adopted conformations were very different than in previously solved crystal structures in cases where these were available (Fig. 3e; Table S3). Binding affinities were determined by global fitting of BLI association and dissociation phases at a range of binder concentrations. There was little correlation between binding affinity and the polarity or intrinsic secondary structure of the target; Kds less than 10 nM were achieved for targets with a wide range of polarities and secondary structure propensities (Fig. 3d-f; Table S4), indicating the generality of the method.

To explore the optimization potential of the designs, we chose a binder, DYNA_1b1, with a Kd of ∼1nM for dynorphin, a kappa opioid receptor (KOR) peptide ligand implicated in chronic pain(*17*, *18*). We used RFdiffusion refinement on top hits as described above for the synthetic targets (Fig. 4a, 2c, and Methods V). Forty-five out of 48 designs showed strong binding in the BLI screening assay at 5 nM, and six had K_d_ <= 100 pM by BLI; fluorescence polarization measurements for two of these optimized designs, DYNA_2b1 and DYNA_2b2, indicated K_d_s <60 pM and <200 pM respectively (Fig. 4b; Fig. S11). Over the set of original and optimized designs for dynorphin A, the peptide populated a wide diversity of random coil, partial strand, and partial helix conformations (Fig. 4c). The dynorphin A binder and dynorphin B binder were orthogonal, binding only to their intended targets (see below), despite having 62% sequence alignment.

**Fig. 4.**
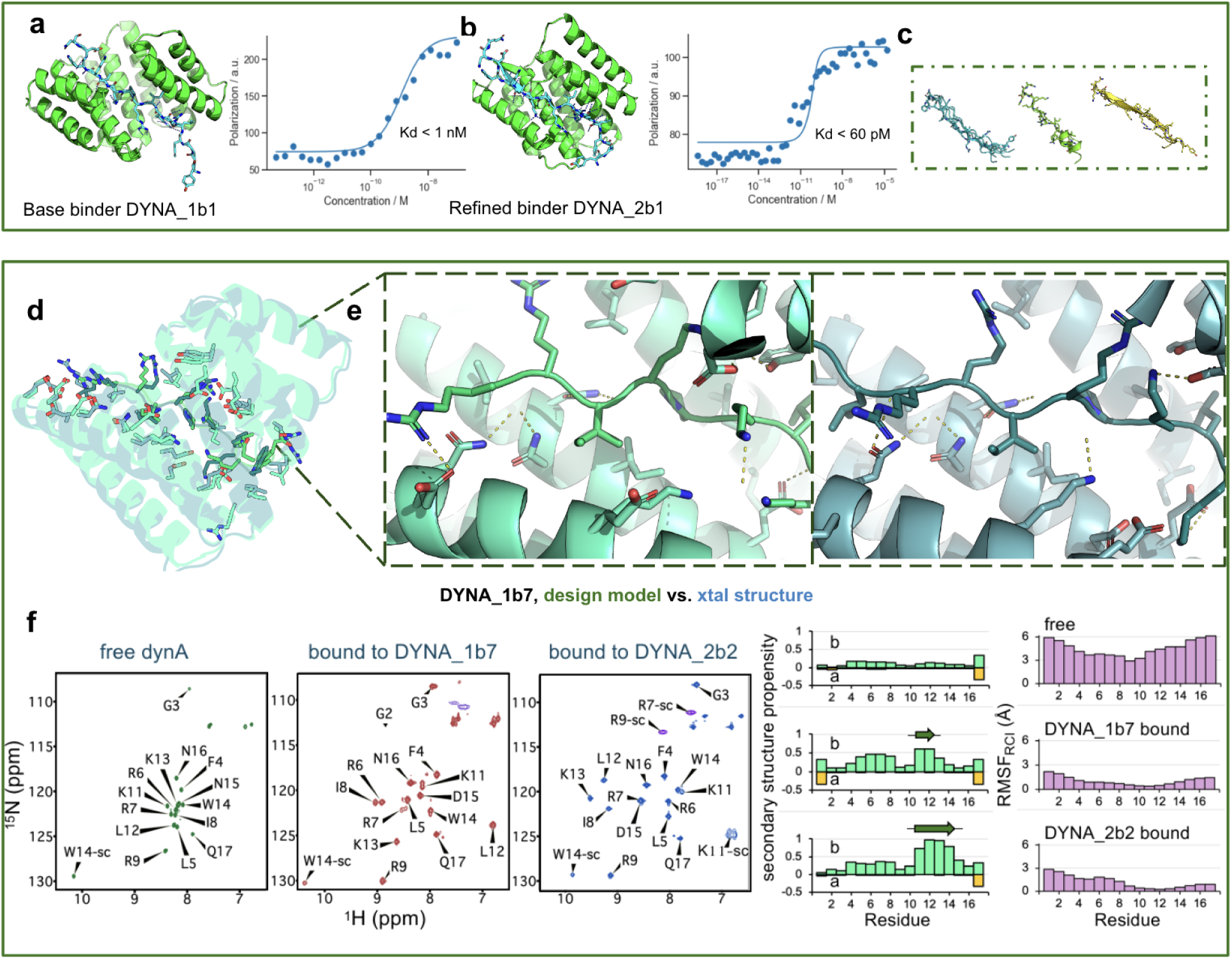
Structural characterization of dynorphin A binder designs. **a,** Design model of DYNA_1b1 bound to dynorphin A in an extended backbone conformation, with five pairs of peptide backbone-protein sidechain bidentate hydrogen bonds (left). Fluorescence polarization (FP) binding with 1 nM TAMRA labeled peptide indicates a K_d_ < 1 nM. **b,** Diffusion refined binder DYNA_2b1 with estimated K_d_ < 60 pM by fluorescence polarization with 100 pM peptide (K_d_ cannot be accurately measured below the concentration of peptide used in the FP assay). **c,** During diffusion-based refinement, the target peptide backbone conformation is sampled as well as that of the binder. **d,** Superposition of the zoom-out computational design model (green) and the 3.15 Å co-crystal structure (cyan) of dynorphin A bound with design DYNA_1b7, with interface residues shown in sticks. **e,** Zoom-in of the design model (green, left) and crystal structure (cyan, right) of the central designed interaction. **f,** Assigned NMR ^1^H-^15^N HSQC spectra (left) of ^15^N^13^C-labeled dynorphin A unbound (free), bound to unlabeled DYNA_1b7, and bound to unlabeled DYNA_2b2 (a variant of DYNA_2b1) in solution, and (right) secondary structure propensity and C*α*RMSF (RMSF_RCI)_) based on backbone chemical shift data. Sidechain Asn and Gln amide resonance peaks in these HSQC spectra are not labeled. When complexed with DYNA_2b2, dynorphin’s Lys11 side chain (sc) amino group, folded from 72.8 ppm in the ^15^N dimension, is observed only because it is stabilized by hydrogen bonding to 2b2 and buried within the complex.

### Structural Validation

We succeeded in solving a co-crystal structure of a 7 nM K_d_ dynorphin A binding design, DYNA_1b7, in complex with dynorphin A (residues 1-17) at 3.15 Å resolution (Fig. 4d-e; Fig. S12). The backbones of both the protein and peptide in the crystal structure match the design model well, with an interface backbone RMSD of 1.2 Å for the complex and interface sidechain RMSD of 2.9 Å (Fig. 4d-e). The key interactions are in the central region of the peptide (Fig. 4e): during design, we excluded the N-terminal YGGF sequence, which is shared between dynorphin A and B and other neuropeptides(*19*), aiming to distinguish closely related peptides in the family where antibodies often fail, and the C-terminal region (-WDNQ) which is missing in some species and hence was not targeted. For this design, the two-sided diffusion refinement eliminated several of the asparagine-peptide backbone bidentate interactions, which could account for the decrease in binding affinity from ∼1 nM for the starting design to 7 nM (Fig. S12; see Fig. S13 for the peptide density maps). The crystal structure confirmed that all the hydrogen bonds in the design model of DYNA_1b7 to the peptide backbone (ASN19, ASN69, ASN70 on binder) were present as designed; the corresponding peptide region (from LEU5 to ARG9) also aligned precisely to the design with Cɑ RMSD=0.6 Å. There were minor shifts of side chains in hydrophobic grooves, and density was missing for the termini that were not included in the design calculations (YGG- and -DNQ).

To investigate changes in dynorphin structure upon binding, we examined the NMR spectra of isotope-labeled dynorphin A unbound in solution, bound to DYNA_1b7 (K_d_ = 7 nM), and bound to the higher affinity design DYNA_2b2 (K_d_ <200 pM; Fig. 4f and Fig. S11). NMR confirmed that free dynorphin A is intrinsically disordered and becomes ordered upon binding except for the regions not included in the design (Fig. 4f). For both bound complexes, the NMR data indicated an extended bound-state conformation, consistent with the design models (Fig. 4f and Fig. S14). The extent of ordering upon binding to DYNA_2b2 was greater than for DYNA_1b7 in both the C-terminal region and around the TRP14-ASN137 bidentate interaction, consistent with the more extensive sidechain-backbone hydrogen bonding in the former (Fig. 4f and Fig. S14).

The extended conformation of the dynorphin peptide in the designed complexes, confirmed by the X-ray and NMR data, is considerably different from any previously solved cryoEM or NMR structures of dynorphin with native KOR (Fig. S14), where it is bound in a compact, partial helix conformation (PDB ID 2n2f, 8f7w)(*20*, *21*). These data highlight the power of computational design for inducing disordered proteins and peptides into non-native conformations.

To assess the contributions of each target residue to binding, we carried out alanine scanning experiments on the dynorphin A binder DYNA_1b1 - dynorphin interaction. In nanoBiT binding experiments, each of nine alanine substitutions on the peptide that disrupt interactions with the binder considerably reduced the extent of binding compared to the wild-type dynorphin peptide (Fig. S15).

### Applications of Designed Binders

Designed binders with high affinity could be useful as enrichment reagents for a broad range of low-abundance human proteins, particularly those involved in signaling pathways. We tested this using as a model the WASH complex, a pentameric complex responsible for the nucleation of branched actin on endosomes(*22*, *23*), and the PER complex involved in circadian clock function. The WASH complex includes WASH (WASHC1), FAM21 (WASHC2), CCDC53 (WASHC3), SWIP (WASHC4), and Strumpellin (WASHC5)(*22*, *23*). FAM21 contains a C-terminal disordered region of about 1,000 residues in length involved in multiple protein:protein interactions (Fig. 5a). We designed three binders for a 27-amino acid disordered window in FAM21 (Fig. 3a). Immunoprecipitation studies showed that FAM21_1b1, retrieved the entire WASH complex from cell lysate (Fig. 5a; Fig. S16). Similarly, a designed binder (PER2_1b1) to a disordered region of PER2 (a phase separating circadian clock protein) enriched endogenous PER2 from cell lysates in pull down experiments (Fig. S17)(*24*, *25*). Designed binders for less well-characterized complexes involving disordered proteins could considerably enhance our understanding of the roles played by this important class of proteins.

**Fig. 5.**
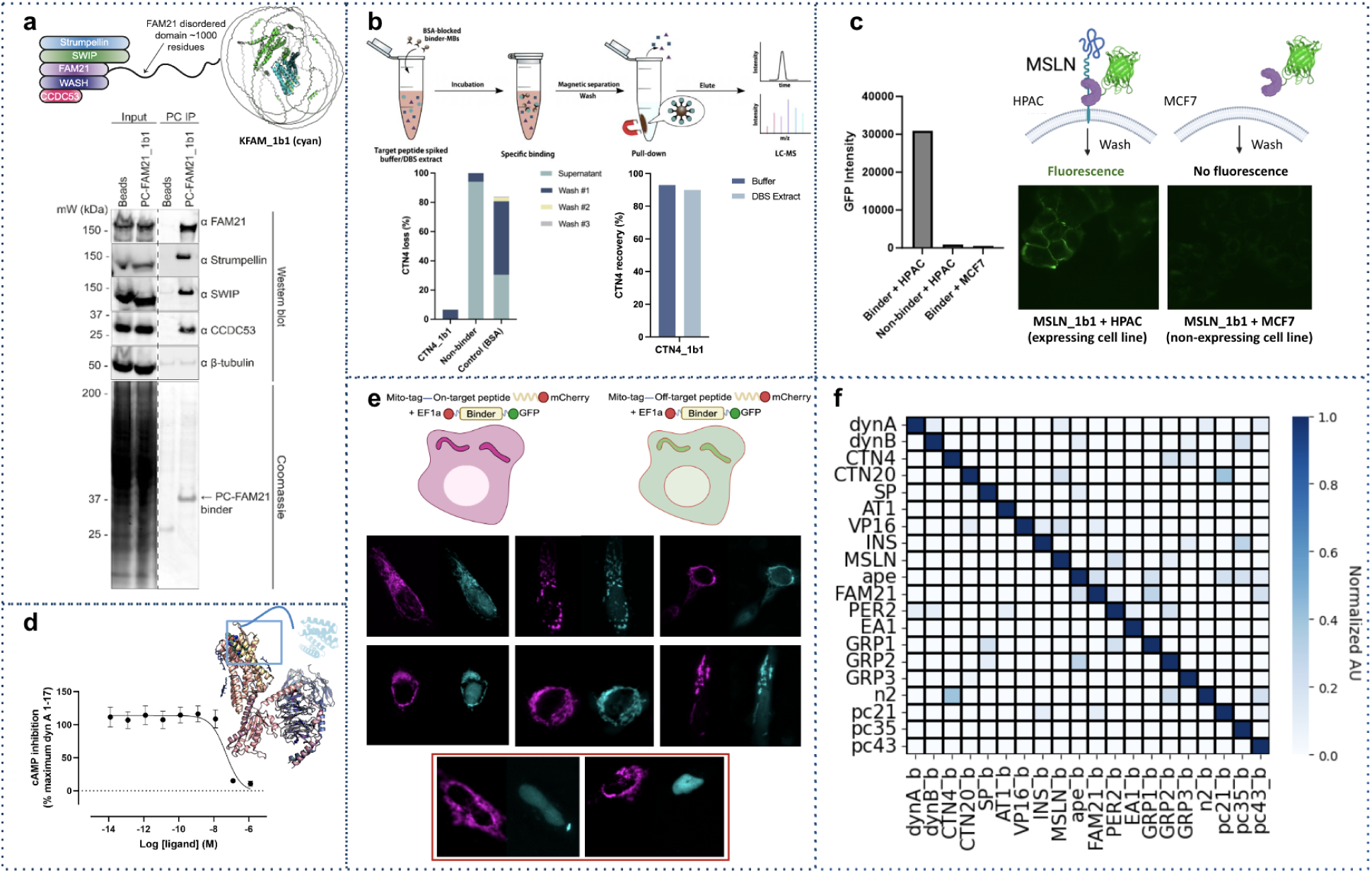
Designed binders are functional and orthogonal. **a,** The WASH complex contains the FAM21 protein with a disordered tail (green cartoon, FAM21; cyan, designed binder). Cell lysates were immunoprecipitated with designed binders, and the bound proteins were assessed by Coomassie stain and Western blot. Designed binder PC-FAM21_1b1 immunoprecipitated FAM21 and other WASH complex subunits from the cell lysate. β-tubulin was blotted as loading control. **b,** Illustration of the use of BSA-blocked designed binder-conjugated magnetic beads (MBs) to capture trypsin-generated target peptides for the case of cystinosin. The amount of peptide recovered from elution was quantified by LC-MS. Left, percentage of unbound peptide from each step normalized to the peak area of peptide standards. Right, the percentage of peptide recovery by LC-MS and normalized to the peak area of peptide standards. Non-binder MBs and BSA-blocked unfunctionalized MBs were used as negative controls. **c**, Designed binder–GFP fusions specifically recognize MSLN targets on cells. MSLN_1b1-GFP staining is observed on the cell surface in MSLN-expressing (HPAC) but not non-MSLN-expressing (MCF7) cell lines following incubation at 1uM. No signals observed when incubating HPAC cells with a non-binder-GFP fusion. **d,** Antagonism of dynorphin A-stimulated KOR signaling by DYNA_2b2 binder competition measured in a cAMP assay in CHO cells. Data are shown as mean ± SEM (n=4). The IC_50_ of DYNA_2b2 binder was 50.3 ± 0.7 nM. See Fig. S19-20 for activation mechanism. **e**, HeLa (CCL-2) cells co-expressing the targets fused to mCherry and a mitochondria-targeting sequence (Mito-Tag) and the binders fused to GFP. GFP signal is only relocalized to the mitochondria for designs with their cognate targets. **f,** 20×20 orthogonality binding matrix determined using BLI. Biotinylated target peptide was loaded onto streptavidin biosensors, and incubated with the designed cognate binder and non-cognate binders. Heat map shows the maximum response signal for each binder-target pair normalized by the maximum response signal of the cognate at 1 uM.

We explored using the designed binders in affinity enrichment coupled with LC-MS for detecting low-abundance peptides generated in proteolytic digests of the proteome (Fig. 5b), such as those from poorly folded mutant versions associated with disease, which can be very difficult to detect by mass spectrometry. To evaluate this, we chose a 12 amino acid tryptic peptide of a mutant form of the lysosomal cystine transporter cystinosin protein implicated in cystinosis, a lysosomal storage disease(*26*, *27*). Binder CTN4_1b1 targeting CTN4 was coupled to magnetic beads and incubated with buffer and blood samples to which CTN4 had been added. LC-MS showed that CTN4 was captured by the binder-conjugated magnetic beads (MBs) but not the control unconjugated or BSA-conjugated beads. CTN4_1b1 enriched and recovered 90% of the CTN4 from both buffer and blood samples (Fig. 5b; Fig. S18; Table S5), a higher recovery than achieved with previously described helical peptide binders(*8*).

Mesothelin is a cell surface glycoprotein upregulated in many cancers that is of considerable interest for tumor targeting (*28*). We investigated whether a designed binder (MSLN_1b1) to a juxtamembrane region of Mesothelin (MSLN) could specifically bind to cells expressing the target (proteolytic cleavage in this region makes more distal regions of the extracellular domain less useful for targeting). We incubated GFP-MSLN_1b1 fusions with cells expressing MSLN (HPAC) and cell lines not expressing MSLN (MCF7), along with a GFP-fusion to a control protein that does not bind MSLN (Fig. 5c). Fluorescence microscopy showed GFP localization of MSLN_1b1 at cell junctions, as expected for MSLN, on HPAC but not MCF7 control cells; no binding to HPAC cells was observed with the control binder (Fig. 5c). Thus MSLN_1b1 specifically binds MSLN on the cell surface.

To date, no antibodies, peptides, or small molecules have been developed to inhibit dynorphin A; existing ligands instead modulate KOR signaling by engaging the deep binding pocket of the receptor(*18*, *20*). To explore the potential of our binders to block KOR signaling mediated by dynorphin A (Fig. S19), we performed an *in vitro* cAMP assay using mammalian cells stably expressing the human KOR. The binder DYNA_2b2 inhibited dynorphin A-dependent KOR signaling with an IC_50_ of 50 nM (Fig. 5d; the IC50 is higher than the Kd because of competition with KOR binding). As noted above, to increase specificity, we excluded the N-terminal YGGF sequence during design to distinguish between dynorphin A and B; this region is critical for opioid receptor activation(*29*) (Fig. S20), and extension to include the YGGF-motif would likely increase potency.

### Binder Orthogonality

We investigated the specificity of the designed binders for 16 native targets and four representative synthetic targets with the most unique target amino acid sequences. The affinities of these designs for their targets are all tighter than 100 nM. We measured the binding affinity for each design against all 20 targets using BLI, and observed little crossreactivity at concentrations up to 1 uM. Within the set of disordered targets considered here, each design thus only binds tightly to the target it was designed to bind (Fig. 5f and Fig. S21).

We investigated the specificity of interactions between the designed binders in cells using a mitochondria colocalization assay(*7*). We expressed six of the designed binders fused to sfGFP(*30*), and the six targeted disordered sequences fused to mCherry and a mitochondrial outer membrane targeting sequence. Each designed binder was expressed with each target one at a time, and binding was evaluated by localization of GFP fluorescence to the mitochondria. We observed localization of the GFP to the mitochondrial for each on-target pair (designed binder with its intended target), but not for any off-target pairs (Fig. 5e), indicating that the designed binders function in cells. As further in cell off-target controls, two sequence homologues of DYNA were also tested and found not to colocalize with the DYNA binder at all (see cartoons in Fig. 5e; and Experimental Methods).

## Discussion

We demonstrate that the conformational heterogeneity of IDPs and proteins can be exploited to make the binder design problem easier than for traditional stable folded targets. For each target, we sampled a wide variety of conformations and identified those compatible with high affinity binding—this contrasts with folded targets, whose fixed structures may admit few optimal binding solutions. Our designs induced the target disordered region to adopt bound structures completely different from those populated in solution or present in previously solved native complexes. Induced fit is a general feature of disordered protein binding interactions in nature(*31–33*), and by taking advantage of induced fit, our approach of threading through a diverse extended scaffold set followed by diffusion-based refinement enables robust computational design of binders to a wide range of disordered sequences, including highly polar, challenging sequences. The design method, which we call *“logos*,” has high computational efficiency and a high experimental success rate (22 designs were tested on average for each case described here), and should be broadly useful for making binders to arbitrary disordered targets of interest (Fig. S22 provides a step by step example of structural components progression of a design campaign). Beyond the examples presented here, the *logos* method has been used to generate binders to Hras, Kras-A, and Kras-B (three polar isoforms out of the four ras isoforms in total), which have high specificity in cells(*34*). More generally, the combination of physically based design approaches for generating a set of privileged starting scaffolds (such as the extended peptide binding protein structures we started with) and deep learning generative methods for introducing diversity and for case-specific refinement could be powerful for many other challenging design problems.

The range of possible applications of designed IDR binders is illustrated by the possible uses of the binders developed here. Cancer and disease-related cell receptors such as Mesothelin (MSLN)(*35*), onco-fusion proteins associated with childhood cancer such as EWS/FLI for ewing sarcoma and EML4-ALK for lung cancer(*36*), and CSP for malaria(*16*) can potentially be targeted and localized through their unique unstructured regions for delivery or degradation. Many transcription factors, epigenetic regulation, and viral-host protein-protein interactions involve intrinsically disordered regions that could be modulated by IDR-binding proteins(*37*). Soluble proteins and peptides ranging from neuropeptides in the brain to poorly studied noncanonical open reading frames (ORFs)(*38*, *39*), which are often intrinsically disordered, such as GREP1(*39*) implicated in breast cancer, could become accessible for imaging and sensing. The potential of our designs to be used in diagnostics is illustrated by the detection of tryptic peptide CTN4 from a mutant form of cystinosin associated with a newborn lysosomal function disorder. More generally, the ability to design binding proteins for short peptide sequences could open the door to next-generation low-cost proteomics platforms based on arrays of specifically designed short peptide binding proteins. For several of these applications, it will be important to extend the characterization of interaction specificity from beyond the sets of targets considered here to the entire proteome. An advantage of targeting long IDRs is that affinity and specificity can, in principle, both be increased if necessary with bispecific constructs targeting two distinct epitopes within the same target.

There are several exciting directions for extending our design approach. First, it should be possible to construct sites appropriate for post-translational modifications such as phosphorylation to enable specific recognition of modified peptides. Second, as catalytic site design methods improve, it should be possible to incorporate proteolytic or covalent modification sites into the designs; the extended conformation of the peptide bond and the pocket-by-pocket sidechain recognition are advantageous properties for protease substrates. The binding pockets and conformations of peptides in most natural proteases resemble our designs, but completely redesigning natural enzyme specificity has proven challenging—instead, designing binders to the target of interest as described here and then incorporating catalytic sites could provide a more general customizable approach.

## Acknowledgments

We thank J.M. Rogers for early discussions of disordered region binding mechanisms, P. Fordyce for inspiring ideas exchange, R.E. Stenkamp believed this was structurally possible. We thank A. Broerman, S. Honda, S. Kenny, and T. Schlichthaerle for valuable conversations to make this work better. We also thank K. VanWormer and L. Goldschmidt for their technical support. This work was funded by The Audacious Project at the Institute for Protein Design (D.H., A.B., C.L., A.K., S.G.), The Open Philanthropy Project Improving Protein Design Fund (H.J., B.C., A.B., S.G., M.A.K.), The Bill and Melinda Gates Foundation, #INV-010680 (A.K., A.K., S.G., X.L.), The Howard Hughes Medical Institute (D.B.), the National Science Foundation Molecular Foundations for Biotechnology No. CHE-2226466 (W.C.), the Defense Advanced Research Projects Agency Harnessing Enzymatic Activity for Lifesaving Remedies program Award HR0011-21-2-0012 (K.W., A.B., X.L.), the National Institutes of Health’s National Institute on Aging U19AG065156 (D.H.), and the Defense Threat Reduction Agency Grant HDTRA1-21-1-0038 (W.Y.). NMR studies were supported by NIH Grant 5R35GM141818 (G.T.M.).

## Author Contributions

K.W. and D.B. conceptualized the *de novo* platform. K.W. developed the computational pipeline *logos*, arbitrary unstructured sequence recognition. H.J. contributed ideas for early-stage di-peptide binder design, contributed the code for threading and parametric perturbation. D.R.H. participated in optimizing the final *logos* pipeline, contributed the code for customized MPNN, customized AF2 (including AF2-multimer, AF2-initial-multimer), and the ideas and code support of local resampling in threading, MPNN-AF2 iteration, organized the github repo. B.C. contributed the code for a customized RFdiffusion build used in the refinement protocol. K.W. with occasional assistance from D.R.H and others, designed, screened, and characterized almost all designs. C.L. helped with some protein purification, and participated in the design to target CSP_N and conducted screening. T.A.R., A.G., and G.T.M. designed, conducted, and analyzed NMR experiments. Y.L. designed, conducted, and analyzed LC-MS experiments in the CTN4 tryptic peptide enrichment detection. K.M. designed and performed the immunoprecipitation assays for the WASH complex, under the supervision of E.D. E.M. designed and performed the in vitro cAMP assay for dynorphin A binders. W.C. offered help and technical support in characterizing some binders. X.L. synthesized and purified peptides, and conducted LC-MS validation of proteins and peptides. S.G., M.Y.L., and A.M. handled large-scale protein purification for crystal sample preparation. A.K.B. and A.K. resolved crystal structures. M.A.K. kindly helped in manuscript writing and figure-making. W.Y. offered help in construct ordering. G.S. under the supervision of S.M.B. offered further downstream characterizations. E.D. contributed conceptual support for the computation idea at the beginning. D.B. supervised the research. K.W. and D.B. drafted the manuscript. All authors reviewed and commented on the manuscript.

### Data and Materials Availability

All data are available in the main text or as supplementary materials. Design scripts and models are available through https://files.ipd.uw.edu/kejiawu/logos.zip and through https://github.com/drhicks/Kejia_peptide_binders/. Crystallographic datasets have been deposited in the Protein Data Bank (PDB) with accession codes 9cce and 9ccf. NMR FID data and resonance assignments have been deposited in BioMagResDB (BMRB) with accession codes 52752, 52755, and 52756.

