## Supplemental Figures for "Design of intrinsically disordered region binding proteins" (from another version of this preprint: version 2, 2024-09-02)

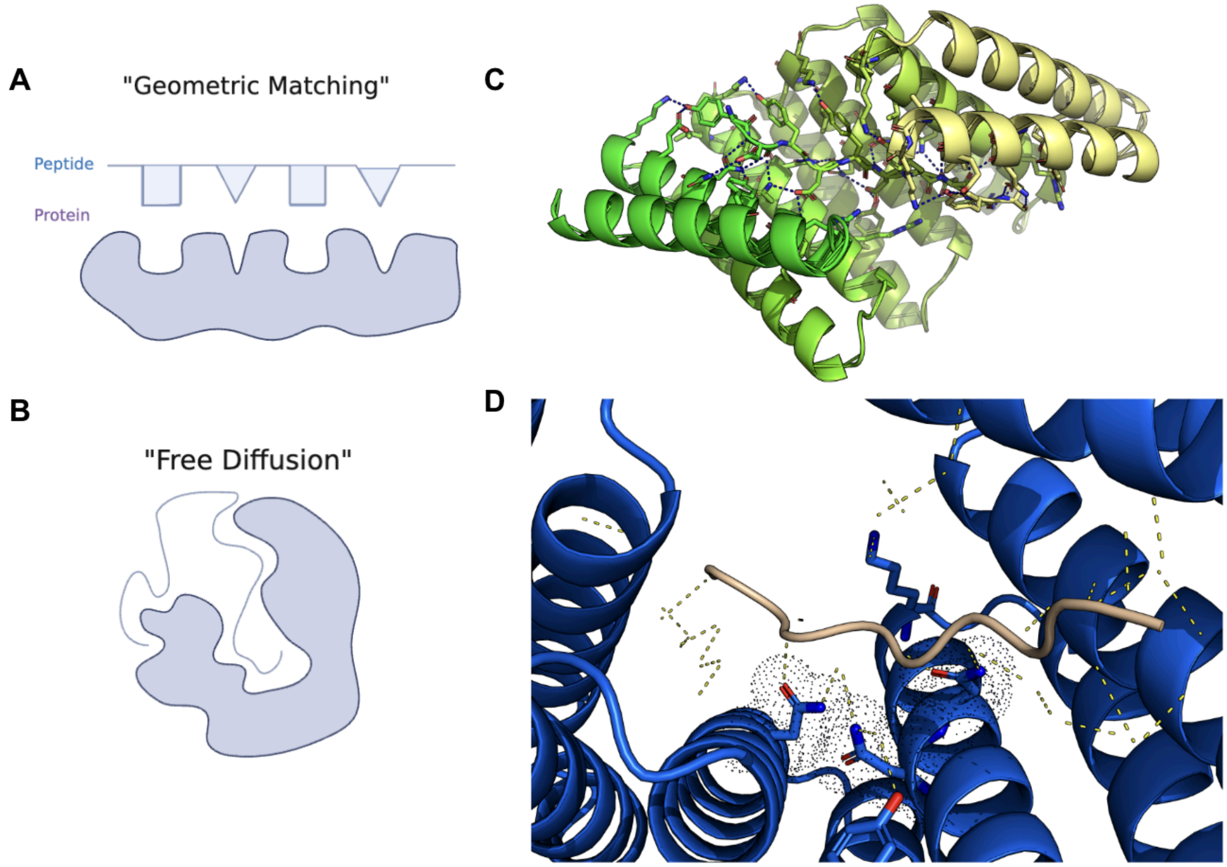


**Extended Data Fig. 1: Template repeat di-peptide binding protein platform.** A) Previous geometric matching approaches required a definite number of pre-built pockets as well as one-to-one parametric matching. B) Free diffusion approach folds relatively large disordered targets (30-40 amino acids) into conformational combinations of alpha-helical or beta-strands, or bind shorter targets in the alpha-helical or beta-strand conformation; C) In the new protein family, a single five-repeat DHR structurally fully surrounds 10 AA peptide target, with adjacent residues on the peptide pointing to the opposite directions to make residue-to-residue interaction (the first residue in each repeat points in one direction, and the second in the other direction to the protein); D) The pre-inserted protein side chain—peptide backbone bidentate hydrogen bonding donors (shown in stick and dot) lock the peptide in more general extended conformations, meanwhile the geometric constraints enable massive protein side chain - peptide side chain hydrogen bonding networks (shown in yellow dotted lines) for affinity and specificity.

| Number of Targets | Optimization | Initial computation | | Computation-experiment cycle | |
|---|---|---|---|---|---|
| | | Base | Diffusion refinement | one-time | two-time |
| 25 | × | ✓ | | ✓ | |
| 2 | × | ✓ | | | ✓ |
| 11 | × | | ✓ | ✓ | |
| 1 | ✓ | ✓ | ✓ | | ✓ |

**Extended Data Table. 2: Summary of computational and experimental protocols.** To summarize, binders to 25/39 targets have not been experimentally or computationally optimized, and produced straight out of the computer through the base design pipeline. Binders to 2/39 targets have not been experimentally or computationally optimized, and produced straight out of the computer through the base design pipeline, but were experimentally tested in two batches to slightly increase the tested number. Binders to 11/39 targets have not been experimentally or computationally optimized, and produced straight out of the computer through the base design pipeline followed by the diffusion refinement. Binders to 1/39 target (dynorphin A) went through experiment-guided computationally optimized. In the first batch, they were produced straight out of the computer through either the base design pipeline or diffusion-refined pipeline; in the second batch, optimized designs were produced by diffusion refinement on the best experimental hits from batch one.

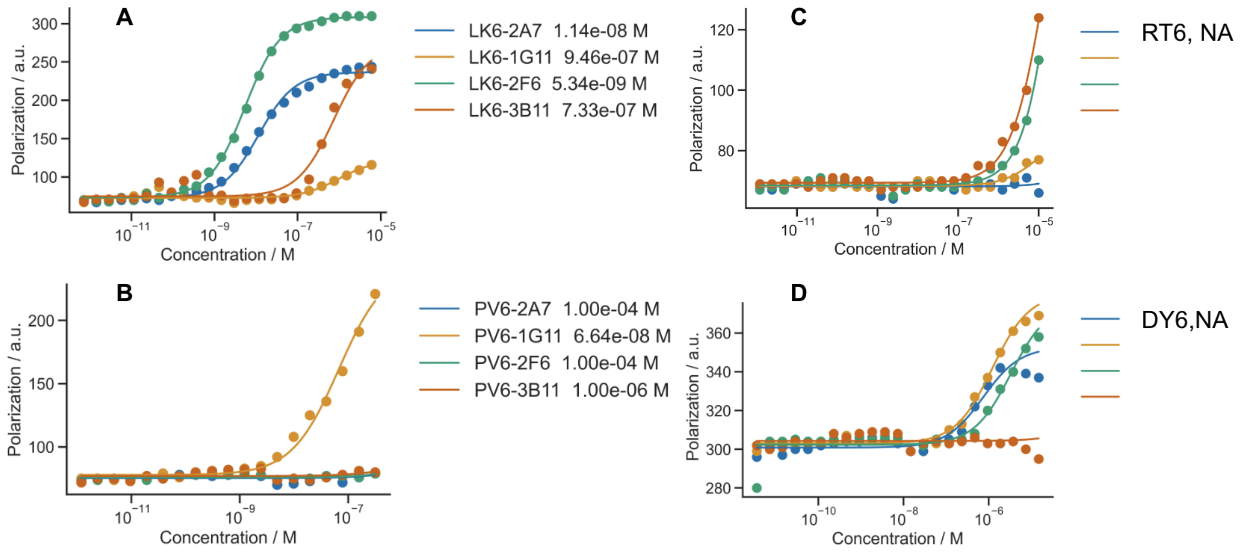


**Extended Data Fig. 3: Initial binding characterizations of the round one di-peptide repeat binders by fluorescence polarization (FP).** FP characterization of the first-round four-repeat designed binders titrating against TAMRA-labeled six-repeat di-peptides. TAMRA-peptides were maintained at 1 nM concentration, while designed binders started from 40 uM (DY), 10 uM (LK, RT), and 1 uM (PV) with two-folded titration.

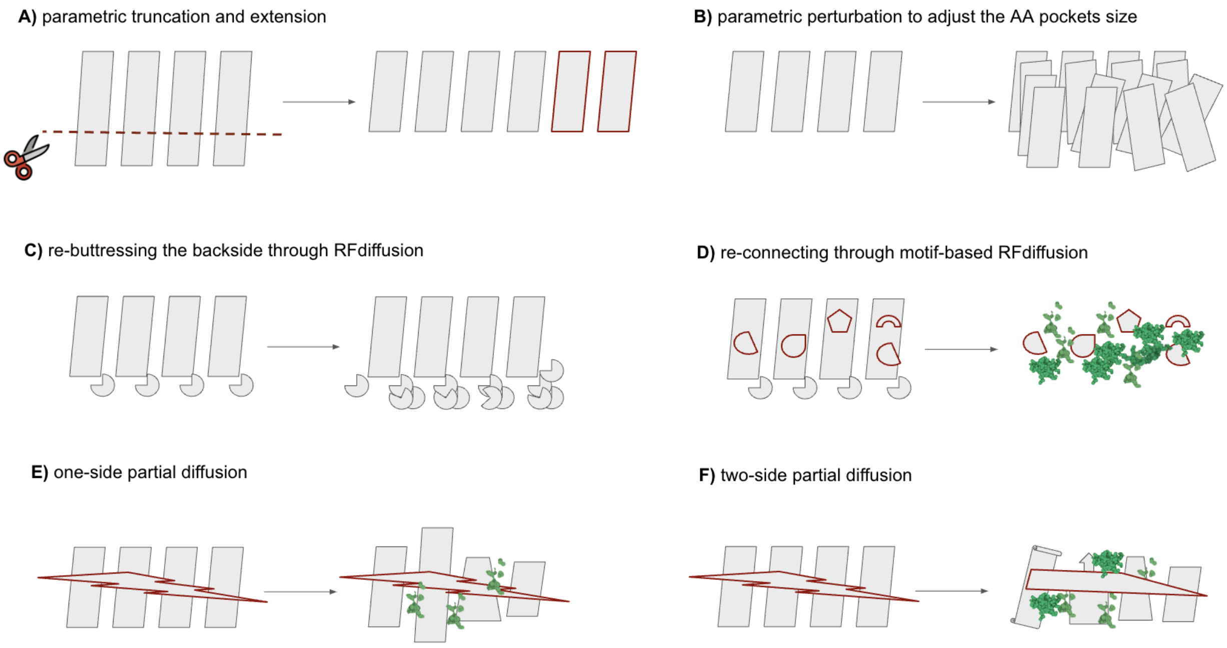


**Extended Data Fig. 4: Template backbone optimization approaches.** Multiple approaches were applied to generate the library templates during pocket design. **A)** Original four-repeat proteins with long helices were truncated by helical length and extended by repeating geometry parametrically to generate five- and six-repeat proteins with short helices for a suitable peptide binding interface. **B)** Parametric perturbations between the repeat-to-repeat transition were applied both symmetrically and asymmetrically to adjust the pocket size. **C)** Each of the original repeat proteins was made of two-helix bundles. To re-buttress and re-pack the core of the protein, all the helices interacting with the peptide were fixed, while the back helices were masked, denoise, and regenerated with RFdiffusion with the context of the target peptide. **D)** Similar to C), per repeat, only the four to nine interacting residues (around the bidentate hydrogen bond donor) to the peptide were fixed. The rest of the protein was masked, denoised, and regenerated with RFdiffusion in the context of the target peptide. **E)** One-side (chain A, which is the designed protein) partial diffusion was conducted as published with PT ranging from [10, 12, 15, 18]. **F)** Two-side (chain A and chain B, which are the designed protein and the peptide target) partial diffusion was conducted with PT ranging from [12, 15, 18, 22].

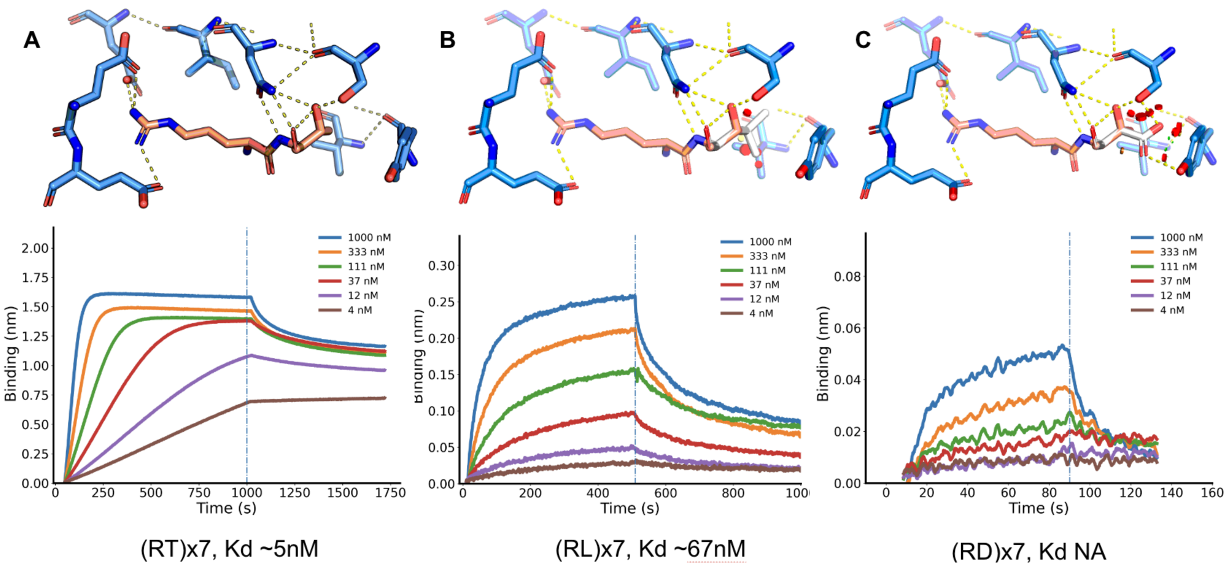


**Extended Data Fig. 5: Repeat di-peptide (RT) binder showed selectivity between threonine, aspartate, and leucine by BLI.** Bio-layer interferometry characterization for the designed (RT) binder titrating against three closely related peptide targets, (RT)x7, (RL)x7, (RD)x7. Three-fold serial dilutions were tested for each binder, and the full tested concentration is labeled. The biotinylated targets were loaded onto the streptavidin (SA) biosensors, and incubated with designed binders in solution to measure association and dissociation.

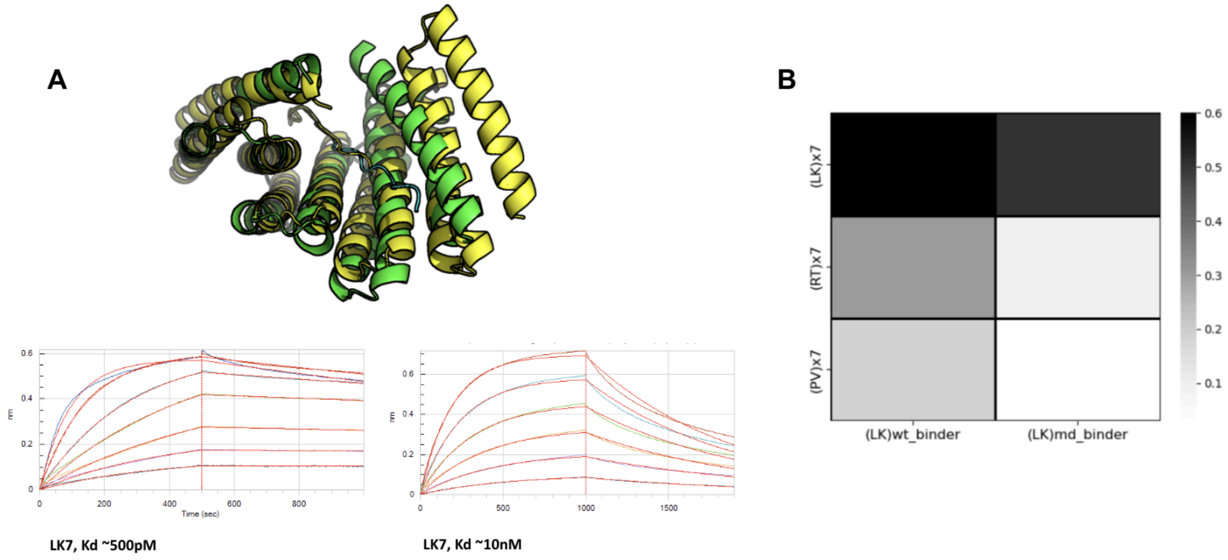


**Extended Data Fig. 6: Refined binder (refinement from motif diffusion) showed increased selectivity while decreased affinity in the case of (LK) binder. A)** WT (LK) binder (cartoon in yellow, BLI plot below, left ) showed a better $K_d$ ~500 pM against its biotinylated target (LK)x7, while motif-diffused (LK) binder (cartoon in green, BLI plot below, right) showed a weaker $K_d$ ~10 nM. **B)** Motif-diffused ((LK)md) binder showed better selectivity against (PV)x7 and (RT)x7 off-targets than WT ((LK)wt) binder in BLI assay. The biotinylated targets were loaded onto the streptavidin (SA) biosensors, and incubated with designed binders in solution to measure association and dissociation. Binders were measured starting at 100 nM concentration, followed by two-fold titrations. Heatmap was plotted with the highest binding signals.

| Base pocket | Related pockets |
|---|---|
| LK | IK,VK, LR, IR, VR, LH, IH, VH |
| RT | KT, HT, RS, KS, HS |
| DY | EY, DW, DF, EW, EF, EY |
| PV | PI, PL |
| GA | GC |
| VP | IP, LP |
| TQ | SQ, SN, TN |
| L | I, V |
| K | R, H |
| R | K, H |
| T | S |
| D | E |
| Y | W, F |
| P | |
| V | I, L |
| G | |
| A | C |
| Q | N |
| M | |

**Extended Data Table 7: Recognition pockets (base and related) generated in this work.** Collection of all pockets generated in this work, with either experimentally validated or computationally predicted confidence in context of disordered targets binding. The related pockets were structurally grouped by their side chain properties, such as charge, polarity, hydrophobicity, aromatics, and size.

A

B

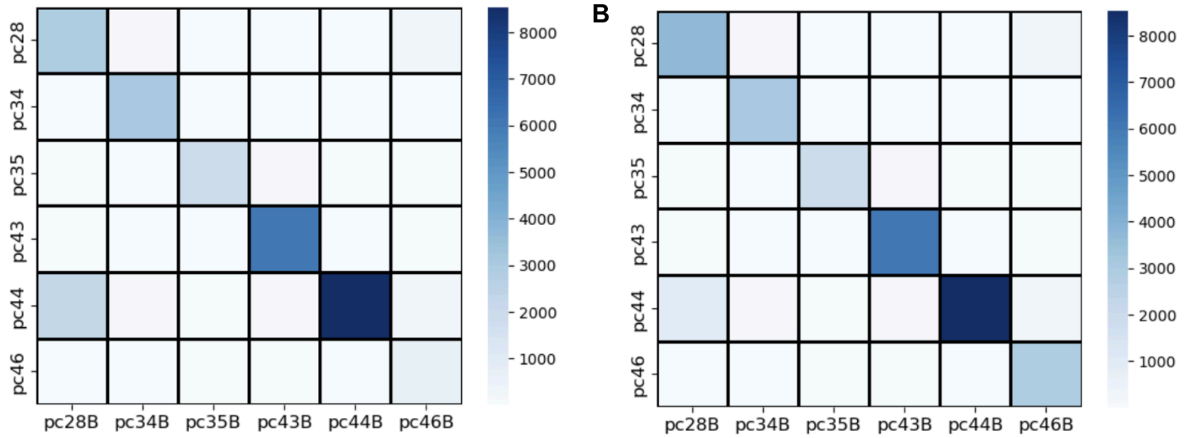


C

| Target | Batch 1 tested designs | Batch 2 tested designs |
|---|---|---|
| pc28 | 10 | 20 |
| pc46 | 7 | 20 |

**Extended Data Fig. 8: Orthogonality improved between targets of arbitrary English words through testing a slightly bigger set of binders (from 8 on average to 20 on average).** Spells out of the English words are as follows: DANIELSILVA (pc28), HIALIENFRIEND (pc34), DERRICKHICKS (pc35), DAVIDLIKESPEPTIDE (pc43), DAVIDITISTIME (pc44), WILLCHEN (pc46). Orthogonality screening was performed by the nanoBiT assay for the six designed binder-synthetic peptide pairs, with lgBiT-binders at 100 nM and smBiT-targets at 0.5 nM. Heatmap indicates the average from N=3 experiments.

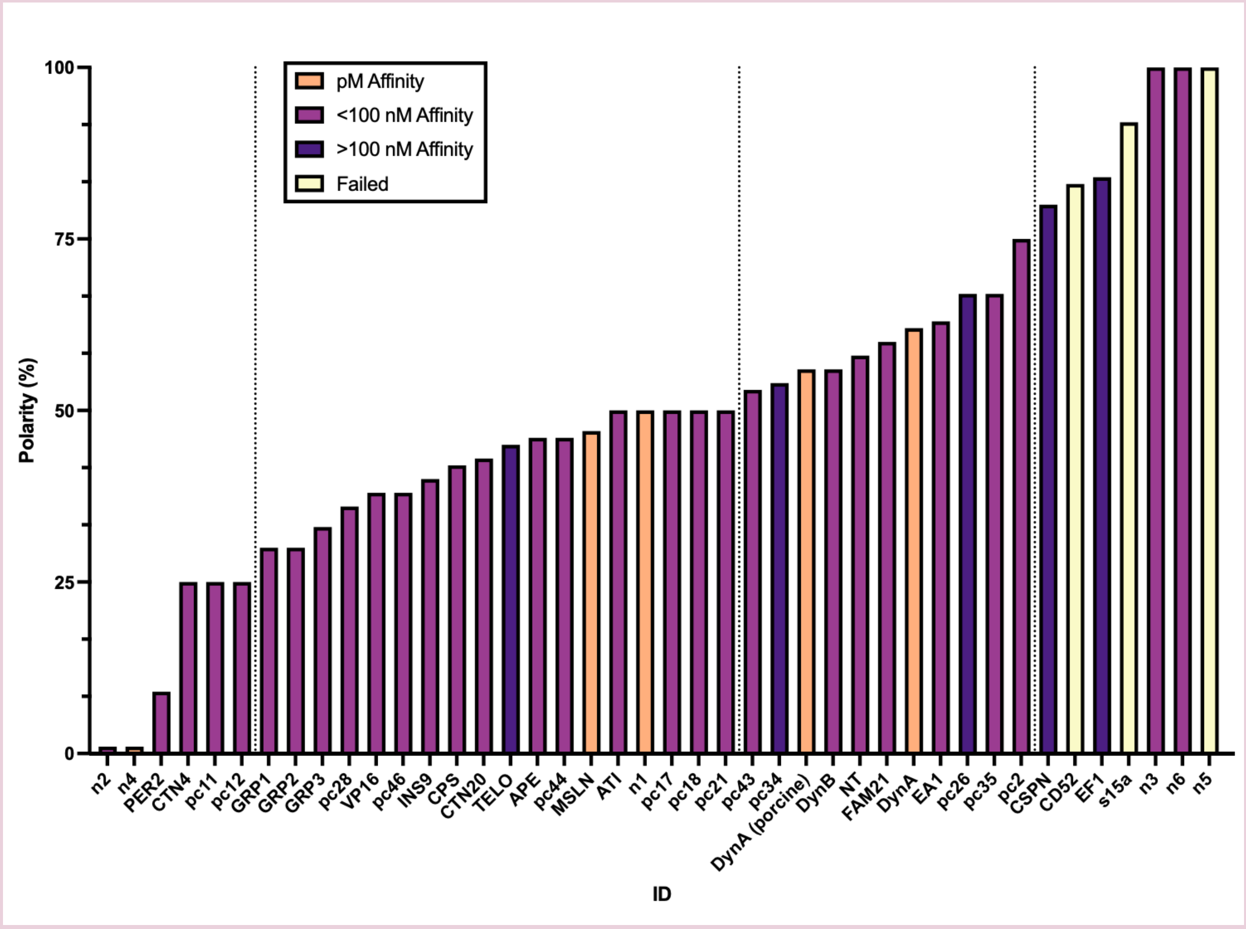


**Extended Data Fig. 9: Target sequence polarity distribution on the 21 native targets and 18 synthetic targets.** Polarity was calculated based on the percentage of polar and charged amino acids (D, E, H, K, N, Q, R, S, T, Y) among the targeted window of each target.

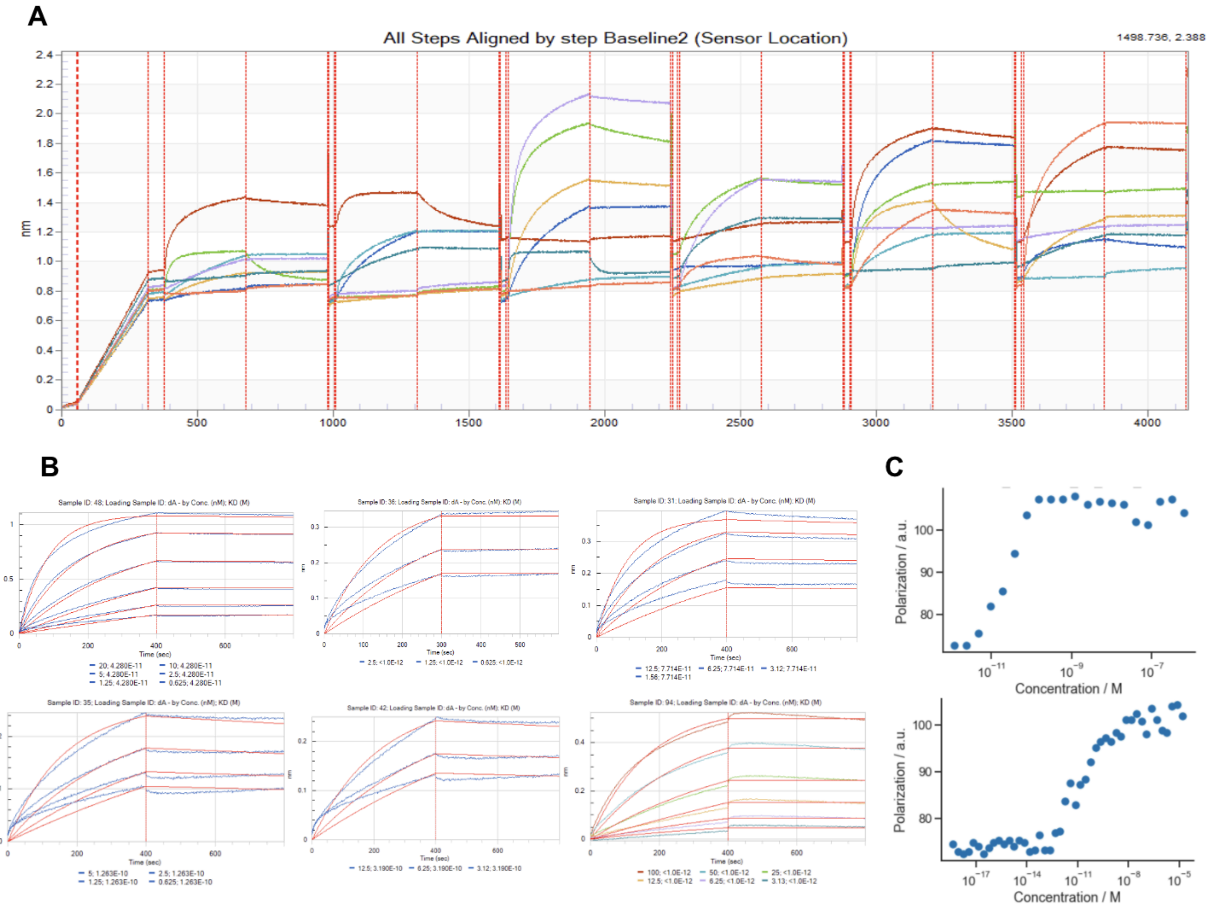


**Extended Data Fig. 10: Tight binding of designed dynorphin A binders.** In the second design round, the optimized dynorphin A binders showed **A)** exceptional screening success rate at 5 nM binder concentration, **B)** ultra-tight binding for six binders' titration by Octet, BLI (tested concentrations and kinetic Kd fittings were labeled under each plot), estimated Kd ranging from <1pM ~ 300pM, and **C)** ultra-tight binding for two binders' thermodynamic fitting by fluorescence polarization (FP) with 100pM TAMRA labeled peptide, estimated as Kd <200 pM, Kd <60 pM (Kd cannot be accurately measured below the concentration of peptide used in the FP assay).

A

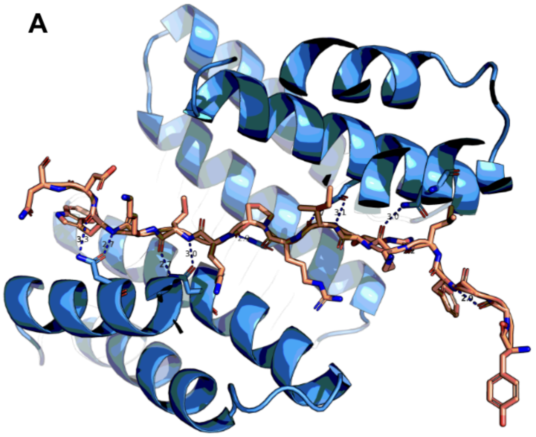

B

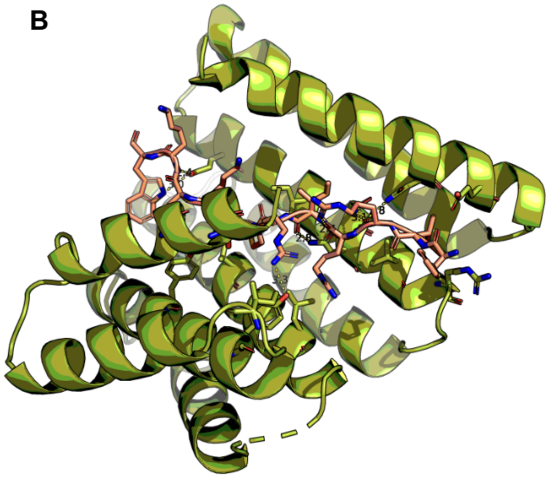

**Extended Data Fig. 11: Comparison of the structure of A) initial design DYNA_1b1 to B) two-sided partial diffusion refined design DYNA_1b7.** During partial diffusion, the binding mode was further changed and diversified on both the binder and the target.

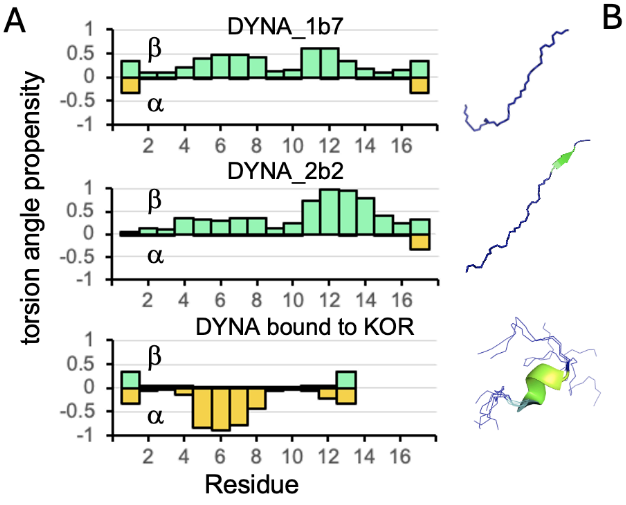


B

| NMR (Cα-Cβ) vs. model comparison | | Predicted by UCBShift from coordinates | | | |
|---|---|---|---|---|---|
| | | random coil | DYNA_1b7 | DYNA_2b2 | DYNA_KOR (2n2f) |
| Measured NMR shifts | free DYNA | **0.02** | 0.36 | 1.01 | 0.52 |
| | DYNA_1b7 | 0.53 | **0.32** | 0.45 | 1.45 |
| | DYNA_2b2 | 0.96 | 0.71 | **0.55** | 1.89 |

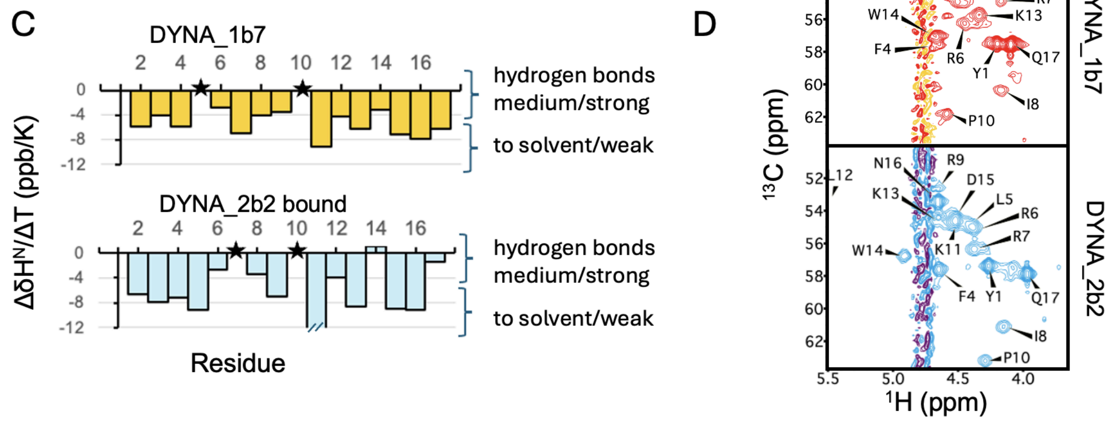

**Extended Data Fig. 12: NMR analysis on dynorphin A complexes.** Dynorphin A binds as extended and partial beta strands as designed. **A)** NMR chemical shift-based analysis using Talos-N of isotope-enriched dynorphins bound to DYNA_1b7 and DYNA_2b2 distinguish helical (α) from extended (β) conformations for each residue along its sequence. Its binding mode is different from the short helix formed when bound to the native human KOR (PDB 2n2f with chemical shifts predicted by UCBShift). **B)** Validation of design DYNA_2b2 by comparison of experimental $^{13}C\alpha$ and $^{13}C\beta$ chemical shifts with back-calculated chemical shifts (scaled $X^2$ analysis) shows that each set of measured NMR chemical shift values agrees best with those predicted from the corresponding atomic coordinates (DYNA_1b7 (PDB 9cce), DYNA_2b2, and PDB 2n2f) by UCBShift (*ref*), rather than with chemical shifts predicted from the other models. **C)** Amide temperature coefficients indicating hydrogen-bond donor amide protons; the backbone amide of W14 of DYNA_2b2 has the strongest hydrogen bond, corresponding to the bidentate hydrogen bonds between W14 backbone atoms and the polar Asn180 side chain of the binder. On either side of W14, there are predicted dynA to binder backbone-backbone hydrogen bonds for K13 and D15 to receptor S5 and K3, in two short antiparallel beta strands; two between K13 NH/CO and receptor S5 CO/NH, and two between D15 NH/CO and receptor K3 CO/NH forming the hydrogen bonded network. The star symbol indicates amide sites not accessible to measurement. **D)** $^{1}H$-$^{13}C$ HSQC spectra of Hα – Cα region. The downfield $^{1}H\alpha$ shifts of L12, K13, and W14 in 2b2 versus 1b7 indicate enhanced binding interactions.

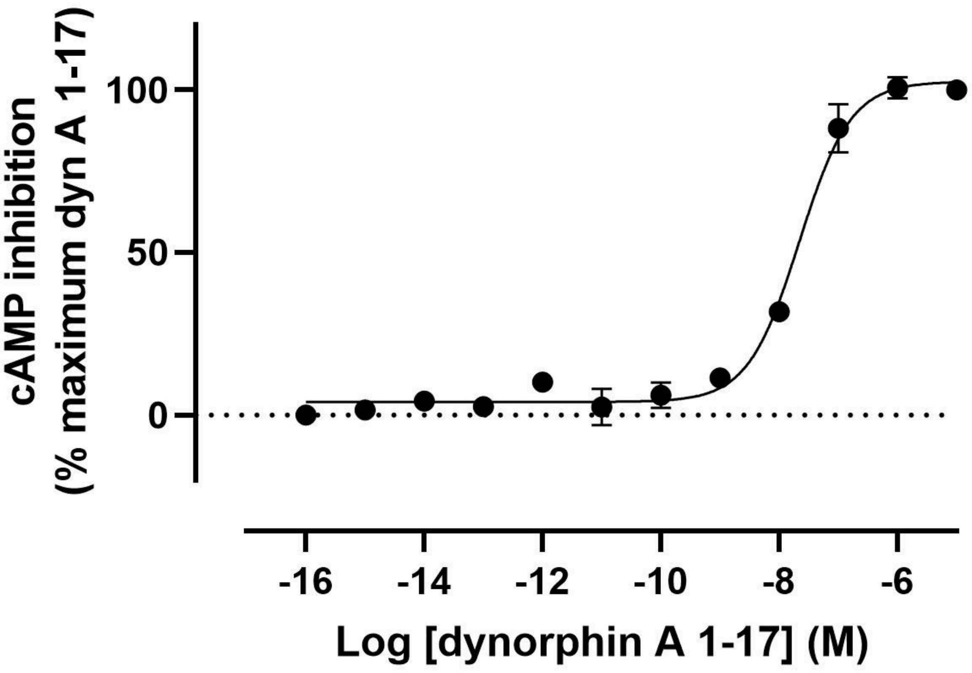


**Extended Data Fig. 13: cAMP assay of dynorphin A (1-17).** Gi-protein mediated cAMP inhibition was measured following KOR activation by dynorphin A (1-17). Dynorphin A (1-17) activates KOR in the cAMP assay with an $EC_{50}$ of 23.1 ± 1.04 nM (mean ± SEM, n=2).

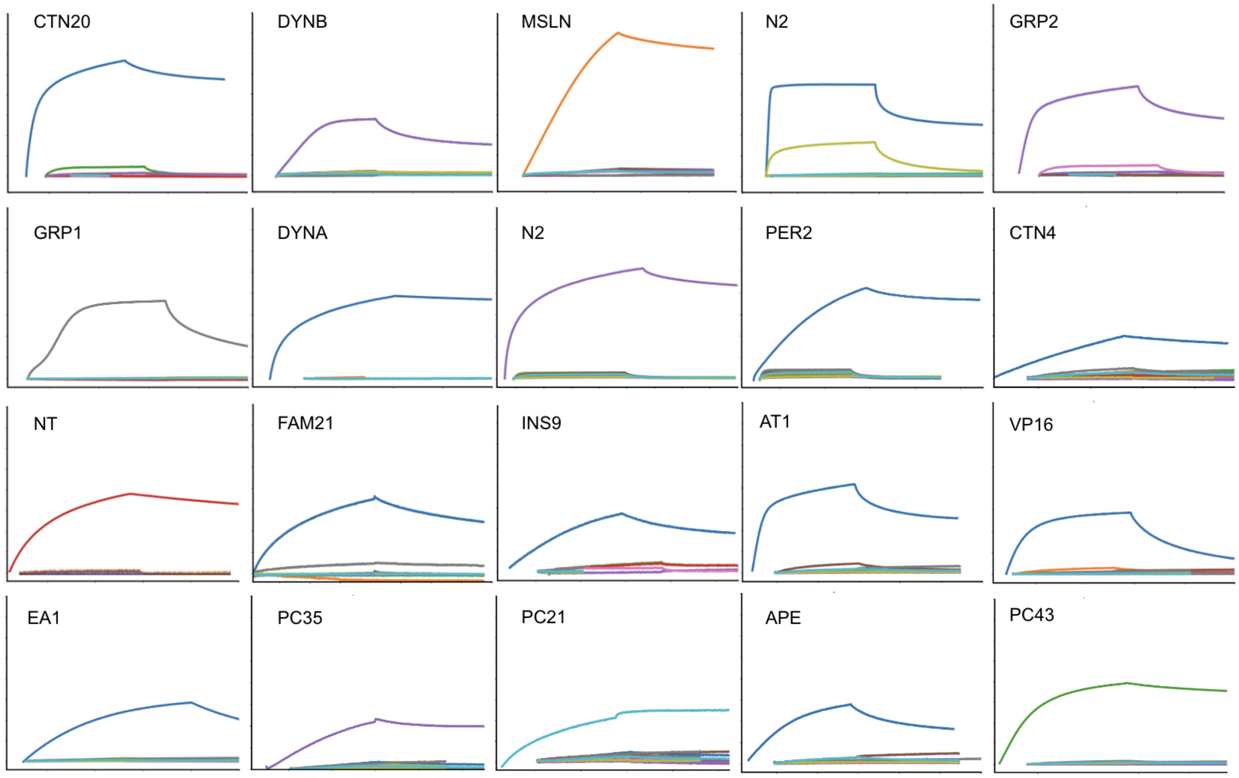


**Extended Data Table 14: BLI specificity test of 20 binder-target pairs with Kd under 100nM.** In each panel, one biotinylated disordered target (as labeled in upper left corner) was loaded onto the streptavidin (SA) biosensors, and incubated with its designed cognate binder and other 19 non-cognate binders all at 1000 nM in solution to measure association and dissociation. Traces were pulled together from multiple individual runs, which have varied association and dissociation times on x-axis. All binding signals were plotted using the same y-axis range (0.0, 2.0 ABU). Among all 400 individual runs, only the cognate pairs were showing obvious strong binding signals; with the exception of target N2 also binds to designed binder CTN4b1 (light green curve), though the non-cognate binding signal showed much smaller with a faster dissociation.

**A**

| ID | Native | Amino acid sequence | Best affinity attained |
|---|---|---|---|
| 1 | DynA | (YGGF)LRRIRPKLKW DNQ | Low pM |
| 2 | DynA (porcine) | (YGGF)LRRIRPKLK | <1nM |
| 3 | DynB | (YGGF)LRRQFKVVT | 9nM |
| 4 | NT | LYENKPRRPYIL | ~1nM |
| 5 | ATI | DRVYIHPFHL | 77nM |
| 6 | APE | QRPRLSHKGPMPA | 92nM |
| 7 | INS9 | SHLVEALYLVCGERG | 59nM |
| 8 | CPS | (SLQP)LALEGSLQ | 100nM |
| 9 | CTN4 | NWLTIFILFPLK | 16nM |
| 10 | CTN20 | ESSVSLTVPPVVK | 10nM |
| 11 | CSPN | DNEKLRKPKHKKLKQ | ~360nM |
| 12 | VP16 | DALDDFDLDML | 18nM |
| 13 | EA1 | PTPGKGPKVYRRKH QE | ~100nM |
| 14 | EF1 | SQQSSSYGQQNPSY DSVRR | >500nM |
| 15 | FAM21 | SSDDDLFQSAKPKPA KKTNPFPLLEDE | 9nM |
| 16 | PER2 | AVPFPAPVPAAY | 67nM |
| 17 | TELO | SWAHPGRTRGPSDR GFCVVSPARPAEEAT SLEGALSGT | 193nM |
| 18 | GRP1 | RRPWVPHLLPFSSPG YLG | 72nM |
| 19 | GRP2 | HENGPWPGPCNARV APMLLPRLPTPGVPS D | 42nM |
| 20 | GRP3 | VLWNSRWPTLQAWG AGLKPGY | ~10nM |
| 21 | MSLN | NGYLVLDLSMQEALS | <1nM |
| 22 | s15a | TSTSPSSSSSSPLSS SSSSSSSSSS | NA |
| 23 | HIS6 | HHHHHH | NA |
| 24 | CD52 | GQNDTSQTSSPS | NA |

**B**

| ID | Synthetic | Amino acid sequence | Best affinity attained |
|---|---|---|---|
| 25 | n1 | LKLKLKLKLKLKLKLK | <500pM |
| 26 | n2 | PVPVPVPVPVPVPVP V | 18nM |
| 27 | n3 | YDYDYDYDYDYDYD YD | 72nM |
| 28 | n4 | GAGAGAGAGAGAGA GA | ~350nM |
| 29 | n6 | RTRTRTRTRTRTRTR T | 5nM |
| 30 | pc2 | LKLKLKLKYDYDYDY D | 84nM |
| 31 | pc11 | LKLKLKLKPVPVPVP V | 50nM |
| 32 | pc12 | LKLKLKLKVPVPVPV P | 36nM |
| 33 | pc17 | DYDYDYDYVPVPVP VP | 45nM |
| 34 | pc18 | DYDYDYDYPVPVPV PV | ~10nM |
| 35 | pc21 | LKLKLKPVPVPVDYD YDY | 27nM |
| 36 | pc26 | LKLKGAPVQQYDYD EHRD | >200nM |
| 37 | pc28 | DANIELSILVA | 3nM |
| 38 | pc34 | HIALIENFRIEND | ~180nM |
| 39 | pc35 | DERRICKHICKS | ~90nM |
| 40 | pc43 | DAVIDLIKESPEPTIDE | 9nM |
| 41 | pc44 | DAVIDITISTIME | 35nM |
| 42 | pc46 | WILLCHEN | 37nM |
| 43 | n5 | RDRDRDRDRDRDRD | NA/10+ uM |

**Extended Data Table 15: Amino acid sequences of all 43 targets we have attempted in this work.** The highest affinity attained was noted in the last column each. The four failed targets (affinity labeled as “NA”) are included.

| Computation method | HBD (hydrogen bonding density / 12AA) | CMS (contact molecular surface / 12AA) | DDG (Rosetta binding energy/ 12AA) | BUNS (buried unsatisfied hydrogen bonds/ 12AA) |
|---|---|---|---|---|
| *logos* | 18 | 620 | -57 | 1.5 |
| PPII | 8 | 360 | -36 | 1.4 |

**Extended Data Table 16: Comparison of the Rosetta metrics of binders designed in this work (*logos*) and the previous poly-proline II repeat peptide binders averaging against the ordered designs.** Calculations were done and averaged over all the experimentally tested designs. Due to the variation on target length, all the calculations were normalized to a base complex unit of per 12 amino acids on the target.
